# Protein deuteration via algal amino acids to overcome proton back-exchange for fast-MAS solid-state NMR of large proteins

**DOI:** 10.1101/2024.01.07.574532

**Authors:** Hanna Aucharova, Alexander Klein, Sara Medina Gomez, Benedikt Söldner, Suresh K. Vasa, Rasmus Linser

## Abstract

With perdeuteration, a current standard for solid-state NMR spectroscopy, large proteins suffer from incomplete amide-proton back-exchange. Using a 72 kDa micro-crystalline protein, we show that deuteration exclusively via deuterated amino acids, largely suppressing sidechain protonation, provides spectral resolution comparable to perdeuterated preparations at intermediate spinning frequencies without proton back-exchange obstacles.

---

Protein perdeuteration has evolved as a fundamental technique in current NMR-based structural biology. Initially introduced for solution NMR, perdeuteration has significantly improved the relaxation properties of ^1^H spins and heteronuclei. Fast magic-angle spinning (MAS) solid-state NMR has emerged as a potent alternative to NMR-based characterization of protein structure, dynamics, and interactions only more recently.^1^ Compared to traditional ^13^C-detected experiments, ^1^H detection has become very popular due to its higher inherent sensitivity. Extensive deuteration of proteins with observation of only a few remaining protons, e. g. on amide-, methyl-or other side-chain sites, has proven to be a viable way to obtain high resolution and sensitivity.^2-5^ Recently, important prospects have arisen from the favorable properties of perdeuterated, micro-crystalline samples in conjunction with increasingly complex pulse sequences.^6-9^

Inasmuch perdeuteration has become indispensable for NMR, it has been imposing severe drawbacks for sample preparation, in particular high costs, low yields, and incompatibility with most potential expression systems. Equally importantly, for efficient amide proton back-exchange, high-molecular-weight proteins with a substantial hydrophobic core usually require unfolding using chemical denaturants followed by renaturation in H^2^O. This is, however, not always feasible, as for many proteins unfolding/refolding results in a major decrease of catalytic activity and/or causes sample loss. Two approaches are usually used to partially evade this bottleneck. One is dispensing deuteration altogether and using ever higher magic-angle spinning frequencies to counteract the line broadening that results from the dense proton network. The smaller rotor diameters required, however, are associated with an impact on the sensitivity of the experiments, such that in our hands the sensitivity afforded with 0.7 mm rotors (ca. 0.5 mg protein) is always significantly lower than what is obtained using 1.3 mm or 1.9 rotors (containing ca 1-2 or 12 mg protein, respectively). In addition, even at 100 kHz MAS, protonated samples have less favorable resonance amide ^1^H linewidths, lower transfer efficiencies, and coherence lifetimes than deuterated samples at 50-60 kHz. For membrane proteins, “inverted fractional deuteration”, *iFD*, is used as an alternative to quantitatively incorporate amide protons while maintaining partial deuteration of the side chains.^10^ This technique involves protonated water while exclusively deuterating the carbon source (usually glucose) of a minimal medium. For membrane proteins, which tend to show slightly broader linewidths compared to micro-crystalline proteins, compatibility of the latter even with larger rotors with their moderate magic-angle-spinning rates (desired to increase the number of spins in the rotor) has been demonstrated. However, due to the rather high remaining proton content (see below), in addition to the impact on coherence lifetimes and transfer efficiencies,^1^ the added homogeneous proton line broadening may compromise the exquisite spectral advantages usually obtained using micro-crystalline samples.

Successful assessment of high-molecular-weight proteins at the frontline of the currently accessible protein complexity is bound to sharp resonances and high sensitivity, generally afforded by sufficient sample amounts of deuterated micro-crystals.^5,6^ As a consequence, any additional line broadening due to added proton content poses a new challenge for the feasibility of conducting a comprehensive NMR characterization of increasingly complex research targets. An approach to harness even higher degrees of residual deuteration while still avoiding the conundrum of proton back exchange, originally suggested for solution NMR studies, is based on perdeuterated amino acid mixes, which can be obtained in an affordable way from algal extracts.^11^ Here we show that using H_2_O with perdeuterated, ^15^N/^13^C-labeled algal extracts for bacterial growth instead of glucose-based minimal media, while circumventing the requirement for proton back-exchange, maintains the very narrow resonance lines of very well-behaved micro-crystalline proteins. The approach is demonstrated to warrant assessment of non-water-exchangeable regions while maintaining favorable sensitivity and resolution for the 72 kDa enzyme tryptophane synthase as a representative high-molecular-weight micro-crystalline protein target with poorly water-accessible hydrophobic core.

The need for proton back-exchange into previously deuterated amide sites in conventional (perdeuteration) approaches is derived from the complete avoidance of protons in D_2_O-based buffers (Fig. 1A). In order to provide the highest deuteration level possible while still affording an expression system based on H_2_O, we turned to fully perdeuterated amino acids, which can be obtained commercially in the form of algal amino acid mixes (not used here) or the whole soluble extracts from algae cultures (used here, Fig. 1B). Such extracts are more often employed as supplements to boost expression in glucose-based minimal media but can also be used as the sole carbon source, e. g. in the case of SILEX supplements to generate labeling schemes facilitating H^α^ detection.^12^ In order to quantify the deuteration levels obtained for the algal extract as a function of sidechain carbon position and compared in particular to the iFD approach, we expressed the SH3 domain of chicken α-spectrin as a well-characterized, low-molecular-weight model system in three different settings and pursued a series of solution NMR assessments. On one hand, we prepared a culture by exclusively mixing H_2_O and commercial ISOGRO powder. In addition, we prepared an iFD sample according to Medeiros-Silva et al.^10^, and finally, a non-deuterated (u-^13^C/^15^N) sample was produced according to standard protocols (see all preparative details in the SI). The samples were assessed in terms of protonation patterns via solution NMR according to previously established principles.^4^ In brief, the H^α^ protonation levels were quantified using a 2D H(N)(CO)CA experiment without proton decoupling in the C^α^ dimension, which allows to quantitatively compare the intensities of proton-coupled C^α^ doublets stemming from CH moieties with the intensities of the CD-derived singlet peaks. Additional ^1^H-decoupled 2D H(N)(CO)CA spectra were recorded to transfer resonance assignments. The protonation of other sidechain carbons was determined using a simple ^13^C constant-time HSQC with both, proton and deuterium decoupling upon indirect evolution, quantitatively comparing the two partially deuterated samples with the non-deuterated one. Figs. 1C and D show the proton content at the H^α^ position, other aliphatic sites, as well the overall protonation of a residue as a function of the amino acid type. Note that not all amino acid types are present or sufficiently dispersed in the SH3 domain, but the general trend is unambiguous.

**Fig. 1:**
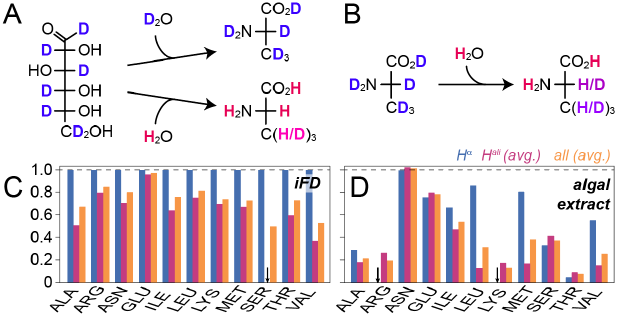
Sidechain deuteration using deuterated algal extracts in comparison to sidechain deuteration from minimal media. **A)** Partial conversion of water protons into fractionally deuterated amino acids in minimal media based on deuterated glucose (top: perdeuteration, bottom: iFD). **B)** Upon employing deuterated amino acids, a high degree of eventual sample deuteration obtained. **C)** Quantitative protonation chart for deuteration based on glucose- and H_2_O-containing minimal medium (iFD labeling). **D)** Protonation chart for water/algal extract-based sidechain deuteration. Amino acids/amino acid mixtures in A/B are generically represented by Ala. The degree of deuteration in A) and B) is tendentially visualized by a blue (deuterated) to red (protonated) color gradient. Black arrows in C) and D) indicate missing peaks, i.e., complete deuteration of those sites, resulting in overall lower protonation ratios of the sample.

In order to test the feasibility of amide-detected experiments without proton back-exchange, we prepared two micro-crystalline samples of *S. typhimurium* tryptophan synthase (TS), a large (2x 72 kDa) α β βα dimer of dimers with a substantial hydrophobic core, according to respective established protocols^6,13^. The first sample was produced in a perdeuterated and back-exchanged fashion, the other using H_2_O and ISOGRO algal amino acids as done for SH3 above. (See preparative and experimental details in the SI.) The samples were assessed in 1.3 mm solid-state NMR rotors, spun at 55 kHz in a 700 MHz magnet at ∼25 °C. (Beyond the sensitivity considerations regarding smaller rotors described in the introduction, larger rotors (1.9 mm) would again either incur broader proton lines and reduced transfer efficiencies or alternatively – in case of a stochastic introduction of amide deuterons – correlations involving more than one amide would be compromised, e. g. through-space correlations or sequential amide-to-amide correlations.^1^) At first, we wondered whether the back-exchange problem known from previous work could in fact be alleviated by the algal amino acid labeling. We hence recorded proton-detected 3D triple-resonance experiments for the perdeuterated and proton-backexchanged protein as well as for the algae-based sample. Fig. 2A shows representative slices from 3D hCONH experiments for the two samples. As hoped for, the additional appearance of correlation peaks is evident. Fig. S2 provides more of such slices for the two samples, in total suggesting that (in the framework of a six-day hCONH experiment for each of the samples, using a threshold of around 20 % relative to the highest-intensity peaks for counting a successful appearance) on the order of 80 additional residues can be found. Importantly, whereas the appearance of many new peaks is obvious, those that are obtained conventionally remain virtually unaltered. To quantitatively address the peak heights of those peaks that are seen in both samples, a correction factor was determined from 1D direct-detection carbon spectra to compensate for slightly differential sample amounts. Fig. S3 shows an overlay of the 1D spectra used for normalization. Fig. 2B shows the normalized peak intensities obtained for the two labeling schemes, whereas Fig. 2C translates the pairwise intensity comparison into a relative intensity difference. Both representations show that the peak intensities of the remaining peaks stay virtually unchanged. A slight increase may be due to peaks that are present even in the perdeuterated samples but have not fully exchanged to 100 % protonation upon sample purification.

**Fig. 2:**
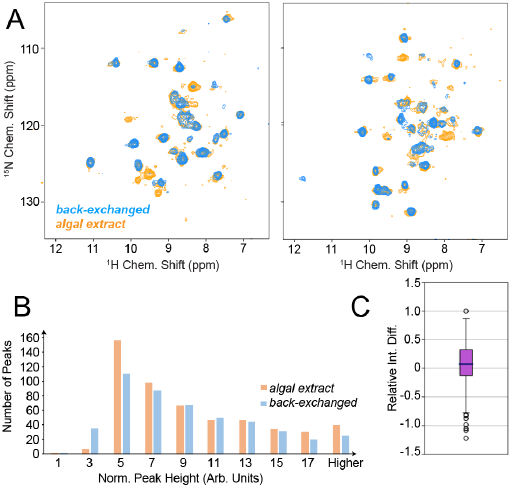
Capturing of water-inaccessible residues in TS through algal amino acid labeling. **A)** Representative slices of a 3D hCONH experiment recorded for TS in either perdeuterated and amide back-exchanged form (blue) or with algal amino acid labeling (orange). **B)** Distribution of peak intensities for peaks found in both of the samples and **C)** the according relative intensity differences for the algal amino acid-based sample compared to the perdeuterated one (i. e., (*I*_algal_-*I*_perdeut._)/*I*_algal_). The bin “5 a. u.” in Fig. 2B corresponds to approximately 20 % of the highest peak intensity found in both samples.

Given the high level of deuteration warranted by algal amino acid labeling in the sidechains (Fig. 1D), one would also expect narrow proton linewidths of algal amino acid samples similar to the perdeuterated/back-exchanged reference case. This is indeed the case: Fig. 3A shows a direct overlay of TS H/N correlations processed via a standard apodization (cos^2^ with shift of the maximum by 4/π). A similar comparison completely without apodization is shown in Fig. S4. Even though most peaks are heavily overlapped in the 2D, the resolved ones visually suggest the linewidth of the perdeuterated sample to be largely maintained upon algal amino acid labeling. To permit a more quantitative comparison, we assessed the proton line widths obtained within the 3D hCONH spectra (Fig. 3B). The comparison of the obtained histograms suggests a virtually identical linewidth distribution. Whereas we estimate the combined uncertainty in measuring the linewidths and determining the mean of the distribution to amount to at least 5 Hz, it becomes obvious that the use of algal amino acid labeling allows to retain the resolution achieved by perdeuteration even for the very well behaved, micro-crystalline sample.

**Fig. 3:**
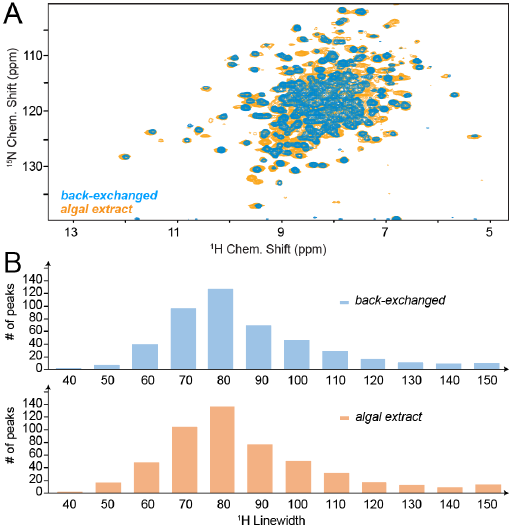
Resolution and ^1^H linewidths obtained upon algal amino acid labeling in TS. **A)** Overlay of H/N correlations for TS obtained at 700 MHz Larmor frequency from a perdeuterated and ^1^H-backexchanged sample (blue) or algal amino acid sample (orange), depicted for an effective proton evolution time of 20 ms and cos^2^ apodization with a shift of the maximum by 4/π. (See Fig. S4 for an overlay without ^1^H apodization.) **B)** Histograms of proton linewidths obtained for perdeuteration/back-exchange (top) and algal amino acid labeling (bottom), assessed in the 3D hCONH spectra. Values in both plots correspond to apodization with 50 Hz exponential line broadening, which value was then subtracted.

The above procedures add a conceptually straightforward and easy-to-implement strategy to obtain well behaved solid-state NMR samples for the highly sought-after 50-60 kHz regime while avoiding refolding for protonation of otherwise inaccessible amide sites. This is of significance in particular for increasingly complex, high-molecular-weight machineries, which have been becoming more and more interesting for recent solid-state NMR studies.^5-8,14,15^ Moreover, the possibility to circumvent not only the requirement for D_2_O-based media but minimal media altogether holds significant promise for solid-state NMR targets with low recombinant expression yields due to toxicity or vulnerability to the conditions of overexpression, including membrane proteins. This consideration is highly protein specific and hence not included as a subject of study here, however, the general trend is clear and has been discussed for solution-state NMR applications previously.^11^ On the other hand, the cost of an exclusively algal amino acid-based culture, prepared as described in the SI, is affordable. (See Table S4 with current price estimates in the SI. Using 10 g of ISOGRO per liter as done here, an increase in costs of ∼1.6 is obtained, however, no systematic optimization was pursued, and less than 10 g of the algal lysate are likely similarly effective.) It is known that part of the amino acid types suffer from acid-or base-catalyzed hydrolysis, which is usually involved for generation of amino acid mixes.^16,17^ This likely explains the relatively high remaining protonation levels in Gln and Glu (Fig. 1D). Rigorous use of (complete) mixes of deuterated amino acid (instead of an algal lysate) might bring the protonation levels of all amino acid types down to the values observed for e. g., Ala, Arg, Lys, and Thr. Solid-state NMR of deuterated proteins, in particular micro-crystalline samples, has the potential to address proteins larger and more complex than those accessible in solution NMR.^6,18^ Additionally, prospects have been emerging for protein sedimentation, albeit with still broader ^1^H lines than found in crystals.19,20 The limitations with respect to residues in the water-inaccessible hydrophobic core has, however, constituted a general, main bottle neck for exploitation of recent technological innovations for high-molecular-weight proteins. This access by amino acid labeling, while maintaining narrow spectral lines, hence provides exciting opportunities for the study of increasingly complex targets. Even if the most insensitive experiments will be compromised by the partial H^α^ protonation remaining, a breadth of experiments will be well feasible from algal amino acid labelled samples that can at least complement the spectra obtained from perdeuterated/back-exchanged samples to fill up remaining gaps in assignment and thus enable comprehensive NMR-based characterization.

In conclusion, we have shown that H^2^O-based expression media that exclusively feature deuterated algal amino acids, which are readily incorporated upon protein expression, instead of more basal carbon sources like glucose or glycerol, while overcoming the barrier of solvent exchange in water-inaccessible protein segments, at the same time largely maintain the spectral properties of perdeuterated samples even under moderate MAS. The approach may help to facilitate comprehensive, high-sensitivity solid-state NMR assessment of deuterated proteins including their hydrophobic core, hence overcoming one of the pertinent bottlenecks for elucidation of increasingly complex biological machineries.

## Acknowledgments

Funded by the Deutsche Forschungsgemeinschaft (DFG, German Research Foundation) under Germany’s Excellence Strategy -EXC 2033 – 390677874 – RESOLV and via the Emmy Noether program. We are very grateful to Prof Dr Leonard Mueller (UC Riverside) for constant advice, protocols, and expressions systems for TS. We are grateful to Prof Dr Paul Schanda (IST Austria) for constructive thought exchange on the topic and sharing their data for a similar scientific endeavour/publication.

## Conflicts of interest

“There are no conflicts to declare”.

## Notes

### Competing Interest Statement

The authors have declared no competing interest.

## References

1. T. Le Marchand, T. Schubeis, M. Bonaccorsi, P. Paluch, D. Lalli, A. J. Pell, L. B. Andreas, K. Jaudzems, J. Stanek and G. Pintacuda, Chem. Rev., 2022, 122, 9943–10018.

2. B. Reif, Chem. Rev., 2021, 122, 10019–10035.

3. R. Linser, M. Dasari, M. Hiller, V. Higman, U. Fink, J.-M. Lopez del Amo, S. Markovic, L. Handel, B. Kessler, P. Schmieder, D. Oesterhelt, H. Oschkinat and B. Reif, Angew. Chem., Int. Ed., 2011, 50, 4508–4512.

4. S. Asami, P. Schmieder and B. Reif, J. Am. Chem. Soc., 2010, 132, 15133–15135.

5. D. F. Gauto, P. Macek, D. Malinverni, H. Fraga, M. Paloni, I. Sučec, A. Hessel, J. P. Bustamante, A. Barducci and P. Schanda, Nat. Commun., 2022, 13, 1927.

6. A. Klein, P. Rovó, V. V. Sakhrani, Y. Wang, J. Holmes, V. Liu, P. Skowronek, L. Kukuk, S. K. Vasa, P. Güntert, L. J. Mueller and R. Linser, Proc. Natl. Acad. Sci. U. S. A., 2022, 119, e2114690119.

7. L. Troussicot, A. Vallet, M. Molin, B. M. Burmann and P. Schanda, J. Am. Chem. Soc., 2023, 145, 10700–10711.

8. H. Singh, S. K. Vasa, H. Jangra, P. Rovó, C. Päslack, C. K. Das, H. Zipse, L. V. Schäfer and R. Linser, J. Am. Chem. Soc., 2019, 141, 19276–19288.

9. S. K. Vasa, H. Singh, P. Rovó and R. Linser, J. Phys. Chem. Lett., 2018, 9, 1307–1311.

10. J. Medeiros-Silva, D. Mance, M. Daniels, S. Jekhmane, K. Houben, M. Baldus and M. Weingarth, Angew. Chem., Int. Ed., 2016, 55, 13606–13610

11. F. Löhr, V. Katsemi, J. Hartleib, U. Günther and H. Rüterjans, J. Biomol. NMR, 2003, 25, 291–311.

12. K. T. Movellan, E. E. Najbauer, S. Pratihar, M. Salvi, K. Giller, S. Becker and L. B. Andreas, J. Biomol. NMR, 2019, 73, 81–91.

13. Y. Tian, L. Chen, D. Niks, J. M. Kaiser, J. Lai, C. M. Rienstra, M. F. Dunn and L. J. Mueller, Phys. Chem. Chem. Phys., 2009, 11, 7078–7086.

14. J. Stanek, T. Schubeis, P. Paluch, P. Güntert, L. B. Andreas and G. Pintacuda, J. Am. Chem. Soc., 2020, 142, 5793–5799.

15. J. S. Retel, A. J. Nieuwkoop, M. Hiller, V. A. Higman, E. Barbet-Massin, J. Stanek, L. B. Andreas, W. T. Franks, B.-J. van Rossum, K. R. Vinothkumar, L. Handel, G. G. de Palma, B. Bardiaux, G. Pintacuda, L. Emsley, W. Kühlbrandt and H. Oschkinat, Nat. Commun., 2017, 8, 2073.

16. R. Linser, V. Gelev, F. Hagn, S. G. Hyberts, H. Arthanari and G. Wagner, J. Am. Chem. Soc., 2014, 136, 11308–11310.

17. D. Schwarz, F. Junge, F. Durst, N. Frölich, B. Schneider, S. Reckel, S. Sobhanifar, V. Dötsch and F. Bernhard, Nat. Protoc., 2007, 2, 2945–2957.

18. H. W. Orton, J. Stanek, T. Schubeis, D. Foucaudeau, C. Ollier, A. W. Draney, T. Le Marchand, D. C. De Paepe, I. C. Felli, R. Pierattelli, S. Hiller, W. Bermel and G. Pintacuda, Angew. Chem., Int. Ed., 2020, 59, 2380–2384.

19. A. Mainz, S. Jehle, B.-J. van Rossum, H. Oschkinat and B. Reif, J. Am. Chem. Soc., 2009, 131, 15968–15969.

20. I. Bertini, C. Luchinat, G. Parigi and E. Ravera, Acc. Chem. Res., 2013, 46, 2059–2069.

